# Romantic jealousy is positively associated with fronto-striatal, insula and limbic responses to angry faces

**DOI:** 10.1101/502096

**Authors:** Xiaoxiao Zheng, Lizhu Luo, Jialin Li, Lei Xu, Feng Zhou, Zhao Gao, Benjamin Becker, Keith M. Kendrick

## Abstract

Romantic jealousy is a complex social emotion combining the different primary emotions of anger, fear and sadness. Previous evidence has suggested the involvement of fronto-striatal dopaminergic circuitry in clinical pathological jealousy, although little is known about overlaps with the neural representation of primary emotions involved in non-morbid jealousy. In the current study, 85 healthy subjects underwent fMRI during resting state and an emotional face recognition paradigm. A total of 150 faces (happy, angry, fearful, sad, neutral) were presented and subjects were required to identify the expression and rate its intensity. Trait romantic jealousy was assessed using the Multidimensional Jealousy Scale. Behavioral results showed that only intensity ratings of angry faces were positively associated with subjects’ jealousy scores. During processing of angry versus neutral expression faces, subjects with higher jealousy scores exhibited greater activation in the right thalamus, insula, fusiform gyrus and hippocampus, left dorsal striatum and superior parietal lobule and bilateral cerebellum and inferior frontal gyrus after controlling for trait aggression and sex. Functional connectivity between the inferior frontal gyrus and caudate was also increased. No associations with resting state functional connectivity were found. Overall, the present study demonstrates an association between romantic jealousy and increased intensity ratings of angry faces as well as in activity and functional connectivity of dorsal striatal-inferior frontal circuitry. Thus, increased emotional responsivity to social threat and enhanced activity in limbic regions and dopaminergic fronto-striatal circuitry may be features of both non-morbid and pathological jealousy.

## 1. Introduction

Jealousy is an important and complex social emotion which can be displayed when an individual is threatened with losing something of personal value and involves affective, behavioral and cognitive components (Harris, 2004; Pfeiffer and Wong, 1989). Although, jealousy is often confounded with envy, it is characterized by distrust, fear of loss, anger, and anxiety, whereas envy is characterized by feelings of inferiority, longing and resentment and disapproval of the emotion (Parrott and Smith, 1993). Jealousy is not considered as a primary emotion in Paul Ekman’s classification, but rather as a combination of the different primary emotions anger, fear and sadness (Ekman, 1999; Hupka, 1984) From an evolutionary perspective, jealousy in a relationship is an evolved adaptation (Buss and Haselton, 2005) that can be beneficial for stabilizing it by providing a warning that a sexual partner is potentially desirable and attractive to others who may therefore compete for them.

Jealousy has mostly been studied in the context of close relationships, particularly romantic jealousy among couples, and is associated with both implicit and explicit self-esteem (Stieger et al., 2012), depression (Aronson and Pines, 1980), autism (Bauminger, 2004) and different romantic attachment styles (Marazziti et al., 2010). Romantic jealousy is comprised of both emotional and sexual components (Weinstein and Wade, 2011) and can occur when one partner perceives a threat of loss of the other to a rival, whether real or imagined. In its most extreme pathological form, romantic jealousy can be delusional and promote aggression in terms of domestic violence, self-mutilation and even murder (Camicioli, 2011).

Initial neuroimaging studies have examined the neural basis of jealousy by monitoring neural reactivity during experimentally induced jealousy. In male monkeys, the amygdala, striatum and superior temporal sulcus (STS), the temporal pole in right hemisphere and bilateral insula are activated when they were confronted with threats to their exclusive sexual access to a female mate (Rilling et al., 2004). Neuroimaging research on pathological jealousy in humans has emphasized the role of dopaminergic fronto-striatal reward circuitry and the ventral medial prefrontal cortex (vmPFC) and insula involved in mentalizing/self-related processing and interoception/salience processing respectively (Marazziti et al., 2013). In line with this conclusion, empirical studies on the behavioral and neural correlates of non-pathological romantic jealousy have reported that the experience of jealousy is accompanied by increased activation in the basal ganglia (BG), and frontal, particularly ventromedial prefrontal (vmPFC), regions and that greater jealousy is associated with an elevated tendency for interpersonal aggression (Harmon-Jones et al., 2009; Sun et al., 2016). Furthermore, there is some evidence for sex differences in neural responses during the experience of jealousy with men demonstrating greater activation than women in brain regions involved in sexual/aggressive behaviors such as amygdala and hypothalamus while women showed greater activation in the posterior superior temporal sulcus (Takahashi et al., 2006). A study of the neural correlates associated with complex social emotions such as envy and gloating has also demonstrated an important role for the vmPFC (Shamay-Tsoory et al., 2007), suggesting that this region is involved more widely in emotions related to jealousy.

Jealousy usually occurs in social interactive contexts, especially relationship triangles, and subtle alterations in the processing of social signals may therefore represent the neural mechanism that promotes its expression. Additionally, it is possible that jealousy may be associated with more general neural processing differences which can be determined even in the absence of external stimulation during resting state conditions. Previous studies in the field have also not controlled for potential overlaps with the neural representation of the individual emotions which comprise jealousy, anger, fear and sadness, and as such we do not know whether there is a neural circuitry which uniquely encodes non-pathological jealousy. An additional issue is to be able to distinguish jealousy from related aggression and both jealousy and aggression can be evoked by a perception of threat (Sun et al., 2016). One way to explore whether altered reactivity towards social signals promotes jealousy as distinct from aggression is to investigate associations between individual variations in trait jealousy and trait aggression and differential responses to social stimuli which convey threat versus neutral or positive social signals. Additionally, associations between trait jealousy as opposed to aggression on neural processing, can be investigated in the absence of external stimuli by analyzing correlations with resting state functional connectivity.

Against this background the current study therefore aimed to investigate associations between trait jealousy and neural activity and functional connectivity using functional magnetic resonance imaging (fMRI) both during the resting state and in response to social emotional signal (face emotion processing) conditions in a cohort of 85 healthy young adults. Given that previous studies have revealed aggressive behavior and sex differences can contribute to jealousy responses (Schützwohl and Koch, 2004; Schützwohl, 2005; Sun et al., 2016; Takahashi et al., 2006) both trait aggression and sex were controlled for during the analyses to account for potential contributions of these factors as confounders. Since jealousy often develops during social contexts, and in interaction with emotional responses of others, we hypothesized that this complex emotion would be specifically associated with neural reactivity towards social emotional signals (affective facial stimuli) rather than the intrinsic interaction of the underlying brain networks during the task-free state. We further hypothesized that trait jealousy would be particularly associated with a heightened behavioral and neural response to social threat signals (i.e. primarily angry and fearful faces).

## 2. Material and Methods

### 2.1 Participants

92 healthy adult Han Chinese subjects (male = 47; age range = 18–27 years, mean age ± SD = 21.68 ± 2.22 years) were enrolled in the present study. 38 subjects (male = 22) were in a current stable relationship and 54 were currently single (male = 25). All volunteers reported no history of medical, neurological or psychiatric disorders, and no history of head injury as well as frequent drug, cigarette or alcohol use and were free of MRI contraindications. All subjects provided written informed consent. The study had full ethical approval by the local ethics committee at the University of Electronic Science and Technology of China and the experiments were carried out in accordance with the latest revision of the Declaration of Helsinki.

### 2.2 Measurements

To exclude potential confounding effects from clinically relevant levels of depression and anxiety, all subjects completed Chinese versions of validated clinical screening scales before the experiment, including the Beck Depression Inventory-Π (BDI-II) (Beck et al., 1996, Wang, et al., 2011) and State-Trait Anxiety Inventory (STAI) (Spielberger et al., 1983; Li and Qian, 1995). Individuals with clinically relevant scores on these two scales were excluded (BDI-II > 28; SAI > 69 or TAI > 69). Individual levels of trait jealousy were assessed using the Multidimensional Jealousy Scale (MJS) (Pfeiffer & Wong, 1989). To control for potential confounding effects of trait aggression, all subjects were additionally administered a Chinese version of the Buss-Perry Aggression Questionnaire (AQ) (Buss and Perry, 1992, Li, et al, 2011).

### 2.3 Experimental procedures and stimuli

Face stimuli from 30 Chinese subjects (15 males) with 5 different emotional expressions (happy, angry, fearful, sad, and neutral) were selected from the Chinese Facial Affective Picture System (Gong, et al., 2011) and the Taiwanese Facial Expression Image Database (TFEID) (Chen and Yen, 2007). All 150 faces were standardized into gray-scale pictures and covered with an oval mask to remove hair and other individual features using Photoshop CS6.0 (see **Figure 1**).

**Figure 1.**
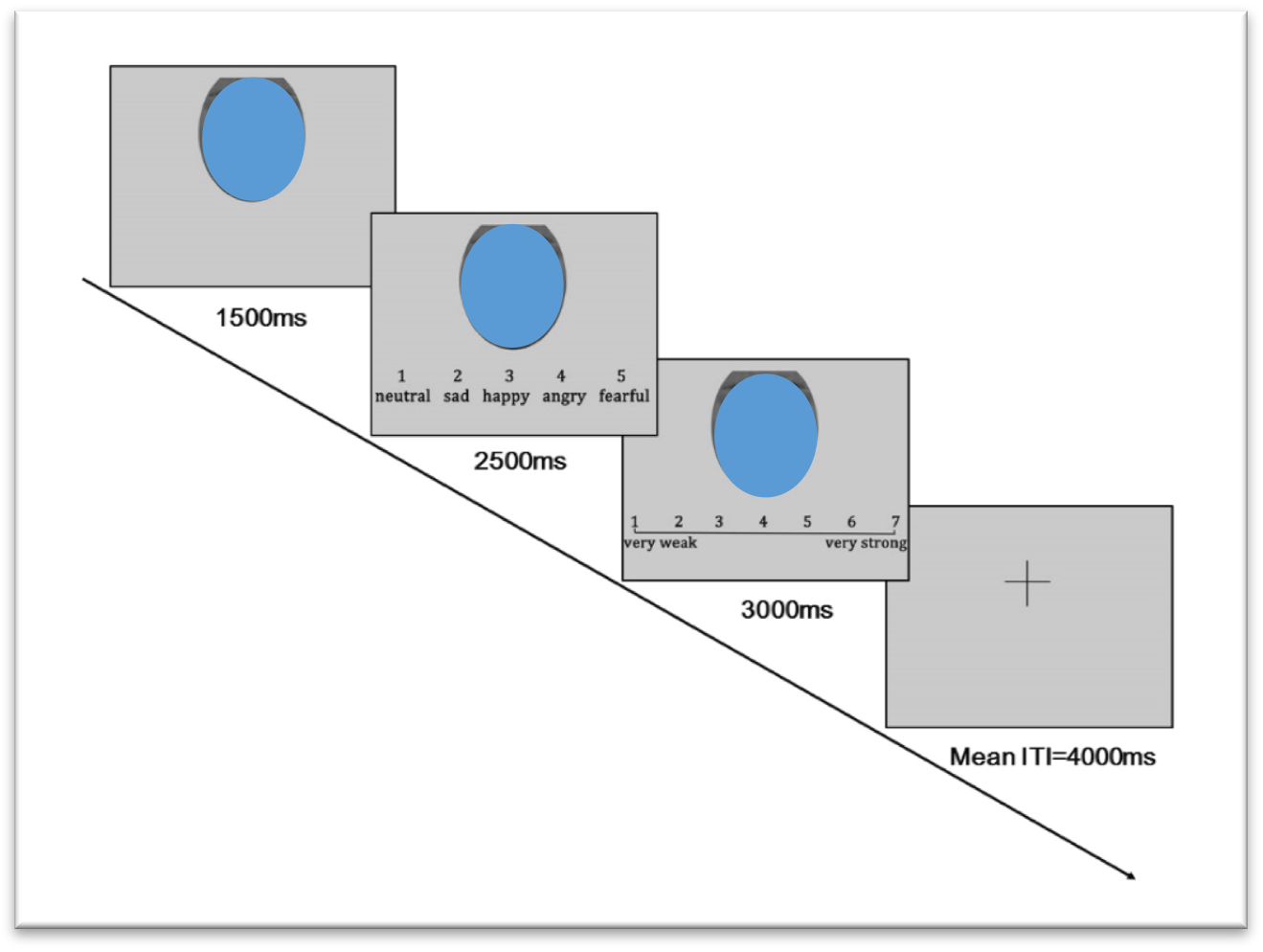
Example trial of the event-related emotional face fMRI paradigm

Before the fMRI session, subjects completed 20 training trials after receiving detailed instructions. Subjects were told to lie still during scanning and foam pads were used to minimize head movement and reduce the impact of scanner noise. The MRI acquisition started with an 8min 30sec resting-state fMRI acquisition. For the resting state acquisition, subjects were instructed to keep their eyes closed and to think of nothing in particular and without falling asleep. Subsequently, subjects performed an event-related fMRI face emotion recognition paradigm. The paradigm consisted of a total of 150 trials that were equally distributed across 3 subsequent runs (50 trials each run, duration 570s per run). Face emotion and gender were balanced across the three runs. Each trial started with passive viewing of the facial stimulus followed by emotion recognition and intensity ratings (total duration, 7000 ms). In the first 1500 ms, a face was displayed for passive viewing. Next, subjects had to indicate the emotion displayed by selecting the corresponding emotion from 5 answer options (1 = neutral, 2 = sad, 3 = happy, 4 = angry, and 5 = fearful) displayed below the face for 2500 ms. Ratings of emotional intensity were subsequently acquired using a separate 7-point rating scale (1 = very weak to 7 = very strong) presented for 3000 ms. A jittered fixation-cross was displayed between the trials for 3600–4400 ms (mean ITI = 4000 ms) and served as an implicit baseline for the analysis. The structure of a trial is additionally visualized in **Figure 1**. The Face Recognition Task paradigm was presented via E-prime 2.0 (Psychology Software Tools, USA, http://www.pstnet.com/eprime.cfm).

### 2.4 Image acquisitions

MRI data were obtained using a 3 Tesla GE Discovery MR750 system (General Electric, Milwaukee, WI, USA) located in the neuroimaging center of the University of Electronic Science and Technology of China. During the task-based fMRI acquisition, a time series of volumes was acquired using a T2*-weighted gradient echo-planar imaging pulse sequence (repetition time (TR), 2000 ms; echo time (TE), 30 ms; numbers of slices, 39; thickness, 3.4 mm; spacing, 0.6 mm; field of view (FOV), 240 × 240 mm^2^; flip angle, 90°; matrix size, 64 × 64). Identical sequence parameters were used for the acquisition of the preceding 8min 30sec resting-state fMRI acquisition. Each run of the Face Recognition Task consisted of 285 volumes and each of the resting state scans consisted of 255 volumes. High-resolution whole-brain T1-weighted images were additionally acquired to improve normalization of the functional images (spoiled gradient echo pulse sequence; repetition time (TR), 6 ms; echo time (TE), 1.964 ms; number of slices, 156; thickness,1 mm; FOV = 256 × 256 mm^2^; flip angle = 9°; matrix = 256 × 256).

### 2.5 Behavioral data Analyses, quality control and assessment of collinearity

In an initial step, a reliability analysis was conducted to evaluate the psychometric quality of the MJS and AQ and both demonstrated excellent internal consistencies (Cronbach’s α coefficients: MJS, α = 0.848; AQ, α = 0.921). Independent sample tests further revealed no sex differences for trait jealousy (*t*_83_ = 0.635, *p* = 0.527) or aggression (*t*_83_ = 0.712, *p* = 0.478) and also no effect of current relationship status (MJS - *t*_83_ = −1.165, *p* = 0.247; AQ - *t*_83_ = 0.349, *p* = 0.728).

Next, the normal distribution of MJS and AQ scores as well as the emotional intensity ratings given during the task and recognition accuracy for all emotional face categories was assessed using Shapiro-Wilk’s tests. Results showed that the AQ, emotional intensity ratings of neutral faces and recognition accuracy for all emotional expression faces displayed a non-normal distribution (*p* < 0.05). Associations between normal distributed indices were examined using Pearson correlations and cases where normal distribution was violated corresponding non-parametric tests (Spearman) were employed. Subsequently, associations between the two scales (Spearman correlation analysis) were explored and associations between MJS scores and the behavioral indices of the emotion recognition task (accuracy) (Spearman correlation analysis) were examined. Pearson correlation analyses were therefore implemented to examine associations between MJS scores and intensity ratings of angry, happy, fearful and sad faces and Spearman correlation analysis between MJS scores and intensity ratings of neutral expression. All these behavioral analyses were conducted using SPSS 18 (Armonk, NY: IBM Corp).

Given that collinear regressors in fMRI models might lead to unreliable estimations (Andrade et al., 1999; Mumford et al., 2015), the variance inflation factor (VIF) (for a similar approach see Chau et al.,2017; Ohashi et al., 2017) was additionally examined to investigate collinearity between the regressors. A VIF > 5 is typically considered to indicate problematic collinearity (Mumford et al., 2015; O’Brien, 2007). VIFs in the present study (factors amongst sex, AQ scores, intensity ratings of angry faces and MJS scores; factors amongst sex, AQ scores, intensity ratings of fearful faces and MJS scores; factors amongst sex, AQ scores and MJS scores respectively) were all < 1.153, arguing against problematic collinearity.

A total of 6 subjects were excluded due to excessive head movement during the resting-state acquisition (head motions > 2.5mm) and 1 subject with a clinically significant depression load and elevated trait anxiety (BDI = 41, TAI = 72), leaving a total of 85 subjects (male = 43, age range = 18–27 years, mean age±SD = 21.64 ± 2.176 years) for all further analyses. A total of 4 subjects exhibited head motions > 2.5mm within a run of the face recognition task (two for Run 1, one for Run 3, and one for Runs 2 and 3), and these specific runs were therefore excluded from further analyses in these subjects.

### 2.6 Task fMRI analysis

#### 2.6.1 Data preprocessing

Preprocessing of the fMRI data from the emotion recognition task was performed using SPM12 (Statistical Parametric Mapping, http://www.fil.ion.ucl.ac.uk/spm) implemented in MATLAB. After discarding the first 5 volumes of each functional time series to achieve magnet-steady images, the remaining images were initially realigned to the first image. To facilitate accurate normalization of the functional images, the T1 structural image of each subject was segmented into grey matter (GM), white matter (WM) and cerebrospinal fluid (CSF) and using skull-stripped bias corrected brain images created using ImCalc. The mean functional image of each subject was co-registered to the structural image and subsequently co-registration was applied to all functional images. The functional images were next normalized to MNI space by applying the normalization parameters that were obtained from the structural images. The resolution of the normalized functional images was 3 × 3 × 3 mm (voxel size). Finally, the normalized data were spatially smoothed with a Gaussian kernel of 8 × 8 × 8 mm.

On the first level, condition-specific regressors for the passive viewing phase of the happy, angry, fearful, sad, and neutral faces were modelled as main conditions of interest. Regressors for the emotion recognition (1= neutral, 2=sad, 3=happy, 4=angry, 5=fearful) and intensity ratings (from 1 = very weak to 7 = very strong), as well as for the six movement parameters were additionally included. For the group level analyses 1st-level emotion-specific contrasts for each facial emotion compared with neutral faces were created (angry > neutral, happy > neutral, fear > neutral, sad > neutral).

#### 2.6.2 Task-based fMRI Analysis

Associations between levels of trait jealousy and emotion-specific neural activity were examined using a whole-brain multiple regression analysis as implemented in SPM12. The whole-brain regression served to identify brain regions where activation showed a linear association with MJS scores for the contrasts angry > neutral, happy > neutral, fear > neutral and sad > neutral respectively. All regression models included sex and AQ scores as covariates.

#### 2.6.3 Task-based Functional Connectivity Analysis

To further explore associations between MJS scores and task-related neural activity on the network level, a functional connectivity analysis was employed using a seed-to-whole brain approach. A generalized form of context-dependent psychophysiological interactions (gPPI) (McLaren, et al., 2012) was implemented to model psychophysiological interactions on the individual level. Seed regions were determined on the basis of the significant results from the BOLD level regression analyses. Seed regions were constructed by defining 6mm radius spheres centered at peak coordinates of significant clusters from the BOLD level analysis using MarsBaR (Brett, et al., 2002). Next, associations between MJS scores and the emotion-specific connectivity of these regions were examined using SPM multiple regression models with the contrasts showing significant results from the BOLD level regression analyses. Again, sex and AQ scores were included as covariates.

### 2.7 Resting State fMRI Analysis

#### 2.7.1 Data preprocessing

The resting state fMRI time series were preprocessed using Data Processing Assistant for Resting-state fMRI (DPARSF) (Yan and Zang, 2010). The first 5 volumes were excluded to achieve magnet-steady images and allow the subjects to adapt to the scanning noise. After slice timing correction, the time series were realigned to the first volume to correct for head motion. Data was discarded if the head movement exceeded 2.5 mm of translation or 2.5 degrees of rotation in any direction. The fMRI images were filtered with a temporal band-path of 0.01–0.1 Hz, normalized using DARTEL and resampled to a 3 × 3 × 3 mm voxel-size. Finally, the functional images were smoothed using a Gaussian kernel of 8 mm full-width at half of the maximum value (FWHM). Six motion parameters, white matter, cerebrospinal fluid and global mean signals were regressed out.

#### 2.7.2 Seed-to-whole brain and seed-to-ROI functional connectivity analysis

To explore whether associations between individual variations in jealousy can already be detected in the absence of external stimulation we computed two resting state analyses. Analysis 1 employed a seed-to-whole brain approach to explore whether resting-state functional connectivity at the whole brain level was associated with MJS scores. Seed regions of interest (ROIs) were defined as a sphere with a 6mm radius centered on the peak voxel of significant associations in whole-brain BOLD level analysis. Analysis 2 aimed at directly examining the pathways that showed associations during task-based functional connectivity by employing a seed-to-ROI approach specifically examining the respective pairs of seed-target regions (seed region and significant target-region from the gPPI analysis). Partial correlation on the extracted connectivity indices was subsequently implemented in SPSS18 to calculate the association between resting state functional connectivity strength and MJS scores with sex and AQ scores as covariates.

#### 2.7.3 Thresholding and mapping

For all whole-brain BOLD level and functional connectivity analyses a consistent thresholding was applied with *p* < 0.05 cluster-level FWE correction (according to recent recommendations to control false-positives in cluster-level based correction methods an initial cluster-forming threshold of *p* < .001 was applied to data resampled at 3 × 3 × 3mm, Slotnick, 2017). Brain regions were identified using the Automated Anatomic Labelling (AAL) atlas (Tzourio-Mazoyer et al., 2002) as implemented in the WFU Pick Atlas (School of Medicine, Winston-Salem, North Carolina).

## 3. Results

### 3.1 Behavioral results

Recognition accuracy for all emotional face categories was high (happy, M ± SD = 98.00% ± 3.40%; angry, 91.30% ± 8.56%; fearful, 88.37% ± 10.01%; sad, 93.80% ± 6.32%; neutral, 91.07% ± 11.49%) demonstrating that the subjects both attentively processed the facial stimuli and correctly identified them. Emotional intensity ratings (7-point scale) given by subjects were also higher for all emotional expression faces compared to neutral ones (happy, M ± SD = 5.04 ± 0.89; angry, 5.22 ± 0.78; fearful, 5.23 ± 0.70; sad, 4.86 ± 0.82; neutral, 3.04 ± 1.76). A repeated-measures ANOVA on accuracy scores with sex (male, female) as a between subject factor and emotional expression (happy, angry, fearful, sad, and neutral) as a within subject factor revealed no main effect of sex (*F*(1,83) = 0.732, *p* = 0.395, 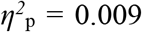) or sex × emotion interaction (*F*(4,332) = 0.624, *p* = 0.611, 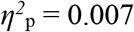). A similar ANOVA for intensity rating scores also showed no main effect of sex (*F*(1,83) = 0.067, *p* = 0.796, 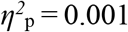) or sex × face emotion interaction (*F*(4,332) = 0.537, *p* = 0.511, 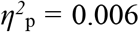). Thus, there were no sex-differences in either recognition accuracy or intensity ratings of the faces.

Correlation analyses showed that MJS scores were positively associated with AQ scores (*Spearman rho* = 0.270, *p* = 0.012), and with intensity ratings of angry (*Pearson’s r* = 0.220, *p* = 0.043) and marginally for sad faces (*Pearson’s r* = 0.212, *p* = 0.052) given by subjects during the task. No significant associations were found between MJS scores and other emotional intensity ratings (all *ps* > 0.121). No significant associations were found between MJS scores and recognition accuracy for all emotional face categories (all *ps* > 0.05). We additionally computed correlations between intensity ratings of face emotions and AQ scores and none were significant (all *ps* > 0.135).

### 3.2 Task fMRI analysis

Controlling for subject sex and AQ scores as covariates, the MJS scores were significantly positively associated with the activity of right thalamus, insula, hippocampus, fusiform gyrus, left dorsal striatum (putamen and caudate) and superior parietal lobule and bilateral cerebellum and inferior frontal gyrus during processing angry relative to neutral faces, and positively associated with superior parietal lobule activation in response to fearful relative to neutral faces (**Table 1** and **Figure 2**). No significant associations were observed between MJS and other face emotion conditions.

**Figure 2.**
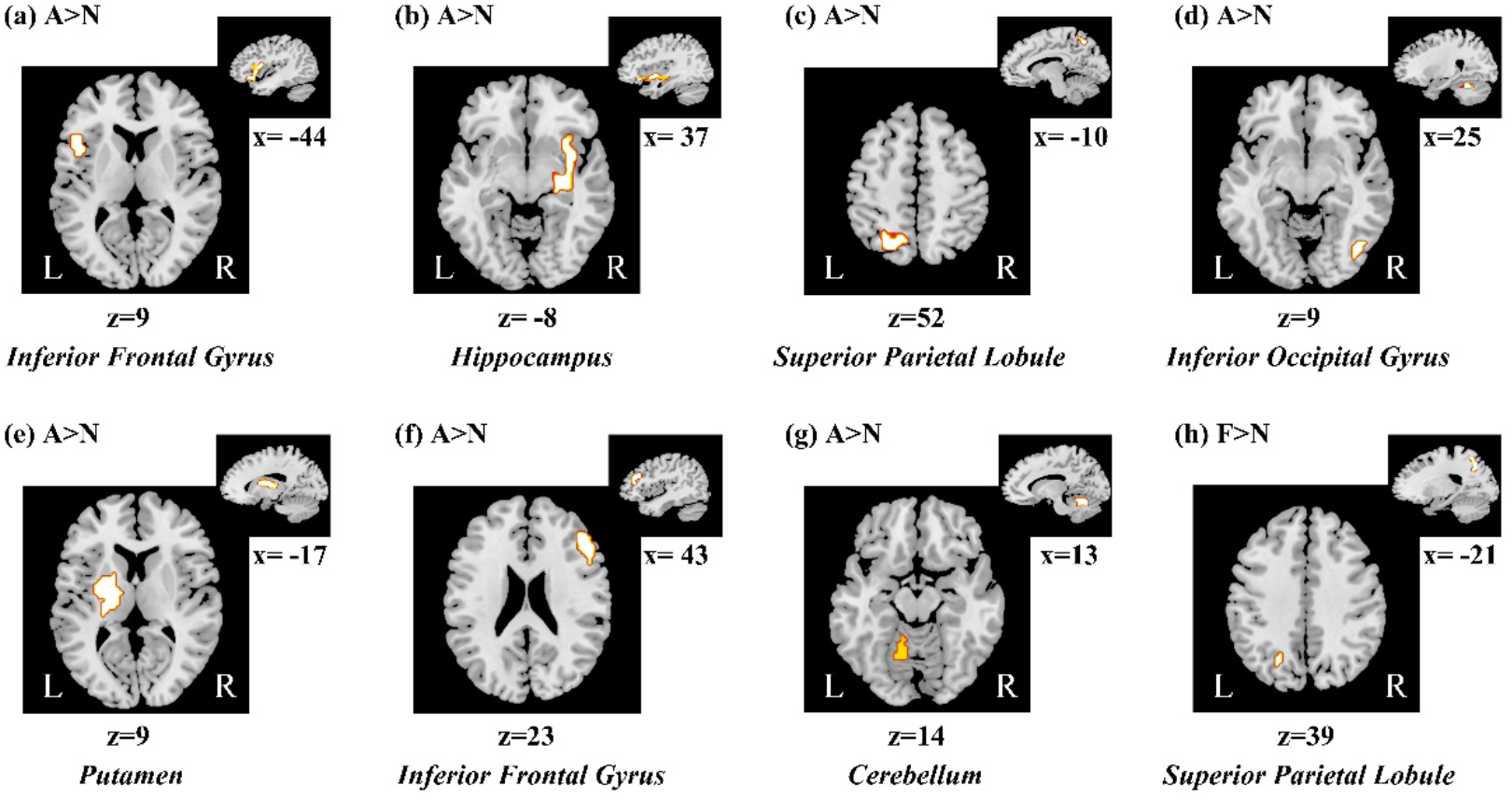
Whole brain analysis. ***(a)****-* ***(g)*** Regions showing increased activation with higher jealousy traits (as assessed by the MJS) during processing angry facial expressions relative to neutral faces. ***(h)*** Regions showing increased activation with higher trait jealousy (as assessed by the MJS) during processing of fearful relative to neutral faces. Both sex and AQ were included as covariates. Findings are displayed at *p* < 0.05 cluster-level FWE correction with a cluster-forming threshold *p* < .001. Coordinates of peak voxels (x/y/z) are given in Montreal Neurological Institute (MNI) space. **Abbreviations:** A>N, contrasts angry > neutral; F>N, contrasts fearful > neutral, L, left; R, right; MJS, Multidimensional Jealousy Scale. AQ, Buss-Perry aggression Questionnaire.

Associations between jealousy and the functional connectivity of these regions using gPPI revealed an association between higher trait jealousy and increased functional connectivity between the right inferior frontal gyrus and left caudate (*k* = 99, *p* = 0.034, *t_80_* = 5.68, x/y/z: -3/17/5) (**Figure 3**) for angry versus neutral faces, with sex and AQ scores being controlled for. No significant associations between MJS scores and functional connectivity were observed during processing of fearful versus neutral faces.

**Figure 3.**
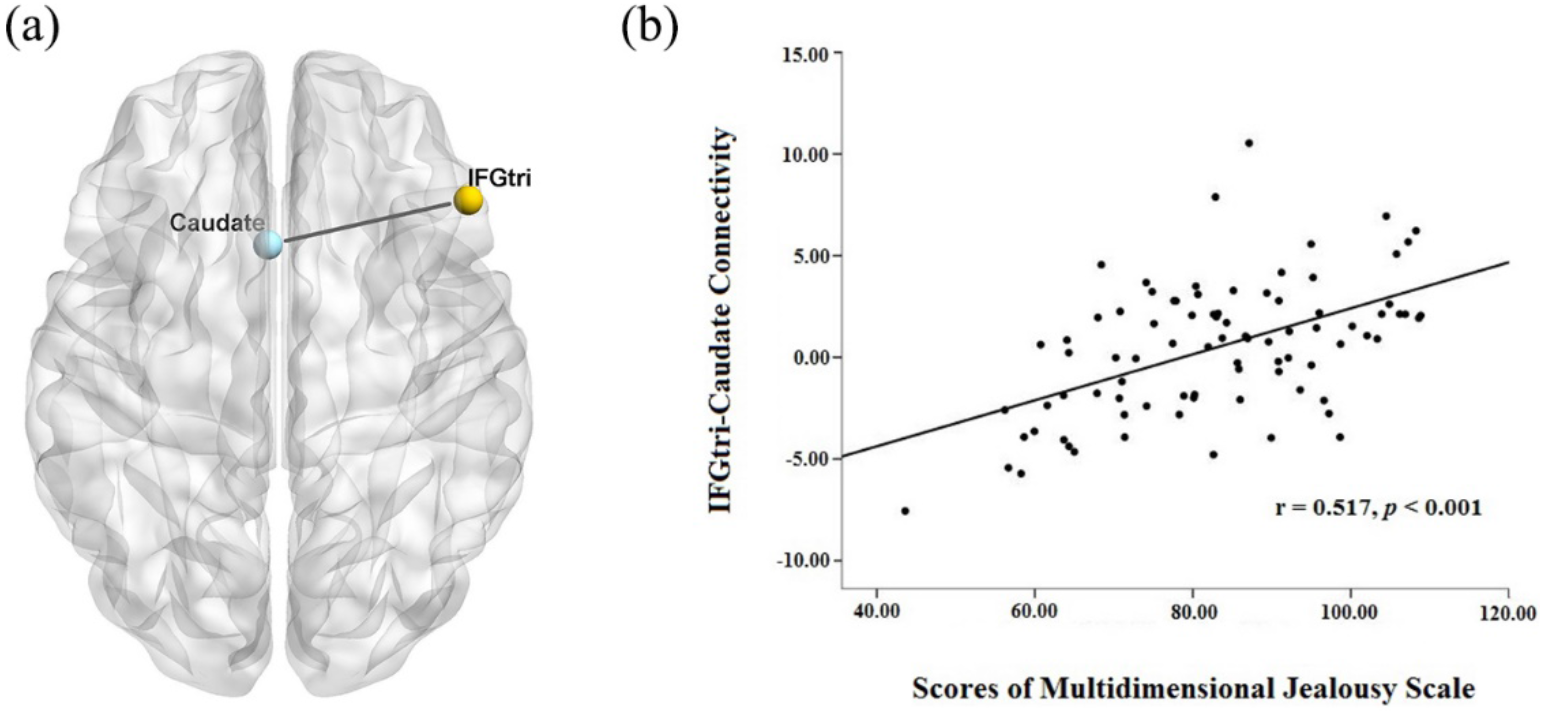
Associations on the task-based network level. **(a)** *Right Triangular Inferior Frontal Gyrus – IFGtri (x/y/z: 51/29/26) as seed region*. Right IFGtri functional connectivity with the left caudate (x/y/z: -3/17/5) was positively correlated with trait jealousy (MJS) during processing of angry relative to neutral faces. **(b)** Scatter plot visualization of the association between trait jealousy and IFGtri-left caudate coupling using extracted parameter estimates. Results were significant at *p* < 0.05 cluster-level FWE correction with an initial cluster-forming threshold *p* < .001. Coordinates of peak voxels (x/y/z) are given in Montreal Neurological Institute space.

**Table 1.**
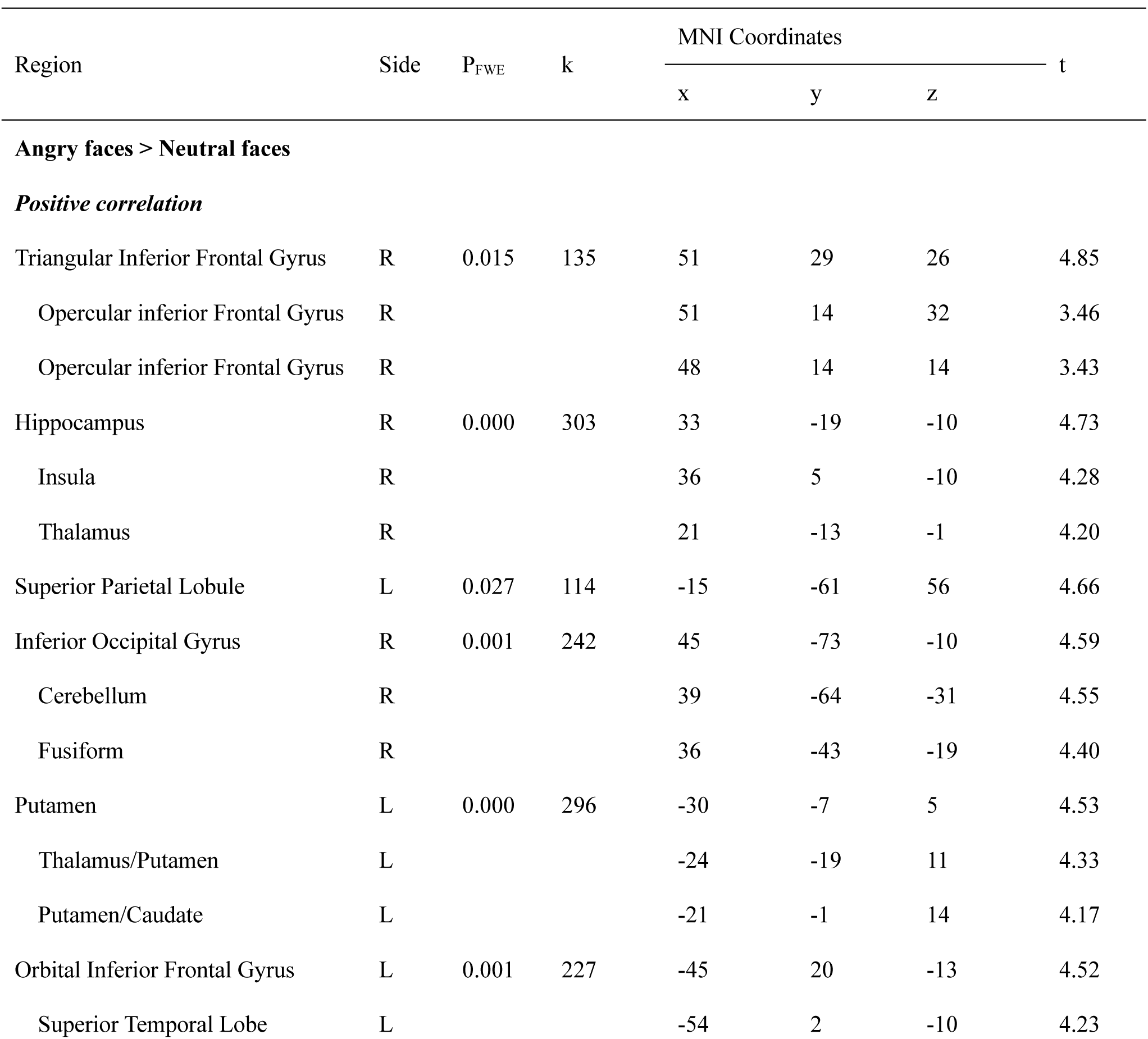
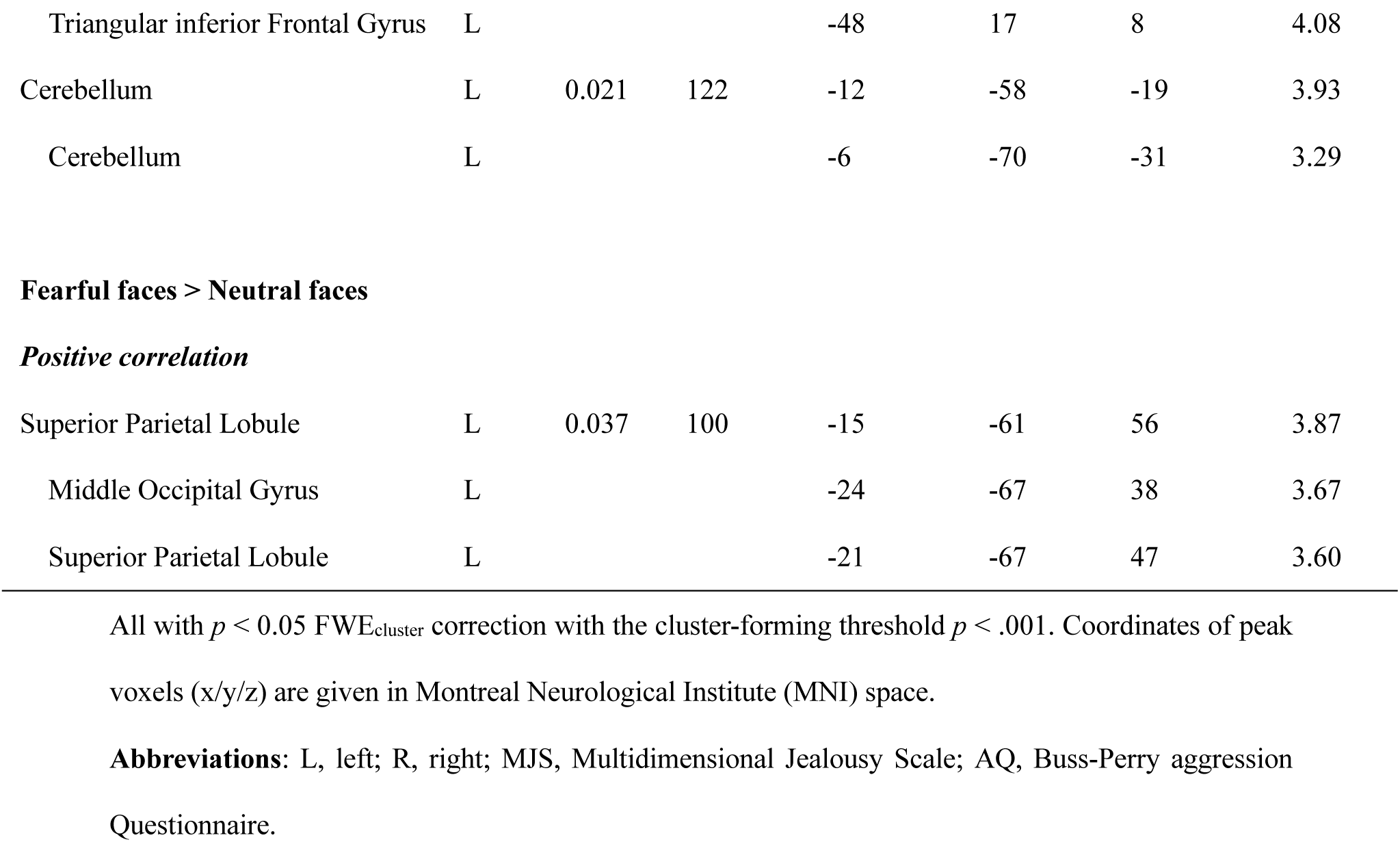
Regions which showed positive correlation with MJS, using multiple regression on whole-brain level with gender and AQ as covariates.

To control for potential effects of co-variations of the intensity ratings with trait jealousy, we re-ran the analyses for the significant BOLD level associations including intensity ratings of angry and fearful faces as nuisance covariates with sex and AQ scores. Results of whole brain analysis and gPPI analysis remained stable for associations with neural responses towards angry faces after additionally controlling for intensity ratings, although not for regions showing correlations with fearful relative to neutral faces (see **supplementary Table S1 and Figures S1 and S2**).

### 3.3 Resting State fMRI Analysis

There were no significant associations between MJS scores and functional connectivity in the seed-to-whole brain analyses of resting state functional connectivity, with sex and AQ scores controlled for. Similarly, there was no significant association between MJS scores and functional connectivity in the seed-to-ROI analysis of the resting state data.

## 4. Discussion

The present study aimed at investigating whether non-pathological levels of romantic jealousy specifically relate to subtle alterations in neural reactivity towards social signals and/or more general alterations in intrinsic neural communication. To this end a dimensional neuroimaging trait approach was employed to determine associations between individual variations in trait jealousy and neural reactivity in response to different emotional facial expressions as well as intrinsic processing in the absence of external stimulation during resting state conditions. Our results revealed that on the behavioral level trait jealousy was specifically associated with increased ratings of the intensity of angry faces. On the neural level, higher levels of trait jealousy were associated with elevated neural responses towards angry relative to neutral faces in the the right thalamus, hippocampus, insula, fusiform gyrus, bilateral cerebellum, IFG and the left dorsal striatum (putamen and caudate) and superior parietal lobule as well as increased functional connectivity between the right IFG and left caudate. The specificity of the neural associations with variations in trait jealousy was established by controlling for trait aggression and sex as well as increased perception of the intensity for angry faces in subjects with higher trait jealousy. Moreover, no associations which survived control for specificity were found in response to other facial emotions or in the absence of external stimuli during the resting state. Together, the present findings suggest that trait jealousy is associated with increased sensitivity to social threat and that, similar to pathological jealousy (Marazziti et al., 2013), it is particularly linked with increased activation and functional connectivity in fronto-striatal circuitry as well as increased activation in limbic and visual processing regions.

Our whole brain analysis of the emotional face task showed that subjects with higher trait jealousy only exhibited stronger neural responses in the thalamus, insula, hippocampus, inferior frontal gyrus, putamen, caudate, fusiform gyrus, visual cortex, superior parietal lobule and cerebellum during processing of angry versus neutral faces. A similar pattern of activation in response to angry faces has been found in previous fMRI studies (see Fusar-Poli et al., 2009), suggesting that the pattern of neural regions responding to angry faces was not influenced by levels of trait jealousy per se but the magnitude of their responses to them. Significant jealousy-related neural responses to fearful versus neutral faces were also found in superior parietal lobe and visual cortex although these were not maintained when intensity ratings were included as a covariant, suggesting that the effects are mediated by differences in the experienced intensity rather than related to trait jealousy per se. In line with this finding, the superior parietal lobe and middle occipital gyrus are both considered as primary sensory regions for visual processing and the middle occipital gyrus is established as being important for perception of stimulus intensity (Cunningham et al., 2004; N’Diaye et al., 2009; Sprengelmeyer and Jentzsch, 2006). Thus the association between jealousy and responses to fearful faces may be driven primarily by enhanced intensity processing, although this is not the case in the context of angry faces where the association with jealousy in these same sensory regions is independent of intensity.

Higher trait jealousy was associated with stronger activation in the bilateral inferior frontal gyrus. Case-studies have reported that lesions in the right frontal gyrus (Saladini et al., 2008) and right orbito-frontal gyrus (Narumoto et al., 2006) are associated with the expression of delusional jealousy. A structural MRI study involving 105 patients also found greater gray matter loss predominantly in the dorsolateral frontal lobes in patients with Othello syndrome compared to matched control patients, indicating that dysfunction of the frontal lobes may be a neuroanatomical correlate for this pathological romantic jealousy syndrome (Graff-Radford et al., 2012). Based on these clinical studies and the involvement of these frontal regions in face emotion processing (Adolphs et al., 1996; Fusar-poli et al., 2009; Nakamura et al., 1999), the stronger activation of bilateral inferior frontal gyrus in the current study might indicate a role in enhanced responsivity to threatening faces in healthy individuals with higher trait jealousy. Additionally, jealousy has been proposed to be an “approach emotion” (Lazarus, 1992) since it is associated with an increased motivation to approach the person towards whom jealousy is expressed. Previous studies have emphasized the role of the left inferior frontal lobe in approach motivation (Gable and Poole, 2014). Thus the association between inferior frontal gyrus activation in response to angry faces and trait jealousy might also reflect a greater approach motivation. Indeed, a previous EEG study on healthy adults reported that evoked jealousy correlated with greater relative left frontal cortical activation in response to a “sexually” desired partner (Harmon-Jones et al., 2009).

The dorsal striatum comprising the putamen and caudate also showed enhanced activation during the processing of angry versus neutral faces in individuals with higher jealousy scores as well as increased functional connectivity with the inferior frontal gyrus. Increased activity in the dorsal striatum has consistently been reported in pathological jealousy (Marazziti et al., 2013). While dorsal striatum activation is associated with the receipt of rewards (Delgado, 2007), it also occurs during the processing of negative valence stimuli (Carretié et al., 2009), including in individuals viewing pictures of someone who has rejected them romantically (Fisher et al., 2010). Thus, it is most likely that greater activation of the dorsal striatum and its functional connectivity with the inferior frontal gyrus reflects an enhanced responsivity to negative emotional stimuli, particularly those associated with social threat. Furthermore, the coupling of basal ganglia and prefrontal cortex had also been demonstrated to play an important role in habit formation (Yin and Knowlton, 2006) and dorsal striatum and related prefrontal connections may be involved in the progressive transformation of jealousy into a habitual behavior (Marazziti et al., 2013).

Fronto-striatal circuitry exhibits a primarily dopaminergic innervation (Björklund and Dunnett, 2007) and dopamine is a key modulator of emotional processes (Sevy et al., 2006). Delusional jealousy is often observed in Parkinson’s disease (PD) patients and several neuroimaging studies have reported that the development of delusional jealousy in PD is significantly associated with dopamine agonist therapy (Poletti et al., 2012), which interferes with reward processing by facilitating dopaminergic bursts and hampering dopaminergic dips (Frank et al., 2004). Indeed, previous neuroimaging studies and patient studies in a number of psychiatric and neurological disorders have generally emphasized the role of dopaminergic fronto-striatal circuits in jealousy (Marazziti et al., 2013). Additionally, increased frontal-striatal activity occurs in obsessive compulsive disorder (Pauls et al., 2014) and obsessional jealousy overlaps with several symptoms observed in disorders with a strong compulsion component.

Higher trait jealousy was also associated with increased insula activity while processing angry compared to neutral faces and increased insula activation has previously been reported in pathological jealousy (Marazziti et al., 2013). Increased insula responsivity in the present results aligns with previous studies establishing its role in the perception and experience of emotion (Kawashima et al., 1999; Phillips et al., 2004; Wicker et al., 2003). Some studies have also suggested that the insula acts as a relay between fronto-parietal regions and limbic regions controlling emotion processing (Carr et al., 2003). Additionally, as a core region in salience network, the insula is specifically sensitive to salient environmental events and facilitates bottom-up access to the brain’s attentional resource (Menon and Uddin, 2010). Thus the insula may contribute to greater jealousy by enhancing the salience of environmental stimuli signaling a potential social threat such as angry faces.

Activation of thalamic and hippocampal limbic regions during processing of angry versus neutral faces was also associated with elevated trait jealousy, and case reports have implicated both in delusional jealousy (see Marazziti et al., 2013). The thalamus controls arousal (Anders et al., 2004; Colibazzi et al., 2010; Etkin et al., 2011; Huguenard and McCormick, 2007), as well as providing a functional link between the frontal cortex and hippocampus (Vertes et al., 2007), and plays an important role in processing visual, auditory and somatosensory information (McCormick and Bal, 1994). Thus, greater activation of these limbic regions in individuals with higher jealousy traits during processing of angry faces may reflect such threatening faces being more emotionally arousing. Interestingly, while the association with trait jealousy was independent of intensity rating scores, when the latter were not controlled for there was a significant association in functional connectivity between the thalamus and IFG. The functional connectivity strength between the IFG and thalamus may therefore play a role in mediating higher ratings of emotional intensity in angry faces in individuals with higher trait jealousy.

The association observed between cerebellar activation in response to angry faces and trait jealousy may also reflect this region’s increasingly recognized role in processing affective stimuli, particularly negative valence ones (Strata, 2015). Indeed, damage to the cerebellum is associated with inability to process negative emotions (Lupo et al., 2015). A case study has also reported delusional behavior in a patient with cerebellar damage (Mitsuhata and Tsukagoshi, 1992).

The absence of any significant associations at the whole brain level between resting state functional connectivity and trait romantic jealousy is perhaps surprising given that some other traits do show changes across healthy and clinical populations (Angelides et al., 2017; Baur et al., 2013; Hahn et al., 2011; Zang et al., 2007). A recent study also reported resting state associations with envy (Xiang et al., 2016), although this was a ReHo analysis rather than functional connectivity per se. Interestingly, this latter study also identified the IFG as showing increased activity in association with dispositional envy and so there may be overlap in frontal regions associated with both jealousy and envy. The lack of associations with resting state indices in the present study may thus indicate that jealousy represents an emotional state which evolves in interaction with social stimuli rather than as an altered intrinsic processing state of the brain per se.

There are several limitations in the present study. Firstly, associations were only made between questionnaire scores for trait romantic jealousy and neural activity in a face emotion task and it would clearly be of interest to investigate if this same circuitry and responses to angry faces is evoked during the actual experience of evoked jealousy. Secondly, while we controlled for a potential contribution of trait aggression on the observed associations with trait jealousy by including it as a covariate, we cannot completely rule out that there may have been some influence of this.

## 5. Conclusions

Overall, the current study explored the neural basis of trait romantic jealousy and the findings provide the first evidence for an association between this and neural responses to angry expression faces and ratings of their intensity in healthy subjects and controlling for trait aggression and sex. Importantly, jealousy-associated activation was found in dopaminergic frontal striatal circuitry associated with pathological jealousy as well as the insula, thalamus, hippocampus and cerebellum which have also been linked with emotional processing and pathological jealousy.

## Acknowledgements

All authors approved the final version of the manuscript. The authors declare no financial interests or potential conflict of interest.

## Author contributions

X.Z., B.B, and K.M.K designed the study and wrote the paper. X.Z. and L.L. prepared all the materials and programed the procedures. X.Z., L.L., J.L., L.X., and Z.G. acquired the data. X.Z., L.L, F.Z., B.B., and K.M.K. analyzed the data. X.Z., B.B and K.M.K interpreted the data and drafted the paper. All authors contributed to and have approved the final version of the manuscript.

