## Supplemental Information for "Romantic jealousy is positively associated with fronto-striatal, insula and limbic responses to angry faces"

**Zheng et al.,**

**Contact:**

### Supplementary Results

**Table S1**

Regions which showed positive correlation with MJS, using multiple regression on whole-brain level with gender, intensity ratings of angry and fearful faces and AQ as covariates.

| Region | Side | $P_{FWE}$ | k | MNI Coordinates x | Y | z | t |
|---|---|---|---|---|---|---|---|
| **Angry faces > Neutral faces** | | | | | | | |
| ***Positive correlation*** | | | | | | | |
| Triangular Inferior Frontal Gyrus | R | 0.013 | 141 | 51 | 29 | 26 | 4.95 |
| Opercular inferior Frontal Gyrus | R | | | 51 | 11 | 32 | 3.51 |
| Superior Parietal Lobule, | L | 0.006 | 168 | -15 | -61 | 56 | 4.73 |
| Middle Parietal Lobule | L | | | -27 | -73 | 32 | 3.73 |
| Superior Parietal Lobule | L | | | -36 | -46 | 56 | 3.46 |
| Hippocampus | R | 0.000 | 242 | 33 | -19 | -10 | 4.72 |
| Insula | R | | | 33 | 8 | -13 | 4.13 |
| Thalamus | R | | | 21 | -10 | -4 | 4.11 |
| Inferior Occipital Gyrus | R | 0.010 | 101 | 42 | -76 | -10 | 4.41 |
| Cerebellum | R | | | 39 | -64 | -31 | 4.35 |
| Fusiform | R | | | 45 | -58 | -19 | 3.61 |
| Putamen | L | 0.000 | 313 | -30 | -7 | 2 | 4.15 |
| Thalamus/Putamen | L | | | -24 | -19 | 11 | 4.35 |
| Thalamus/Putamen | L | | | -18 | -10 | 11 | 4.12 |
| Cerebellum | L | 0.001 | 205 | -15 | -61 | -19 | 4.20 |
| Vermis | L | | | 3 | -58 | -28 | 3.33 |
| Orbital Inferior Frontal Gyrus | L | 0.002 | 169 | -45 | 20 | -13 | 4.19 |
| Opercular Inferior Frontal Gyrus | L | | | -45 | 5 | 14 | 4.15 |
| Superior Temporal Lobe | L | | | -54 | -1 | -10 | 3.96 |
| Fusiform | R | 0.043 | 98 | 36 | -43 | -19 | 4.17 |
| Cerebellum | R | | | 18 | -61 | -19 | 3.76 |
| Cerebellum | R | | | 27 | -49 | -25 | 3.64 |

All with $p < 0.05$ $FWE_{cluster}$ correction with the cluster-forming threshold $p < .001$. Coordinates of peak voxels (x/y/z) are given in Montreal Neurological Institute (MNI) space.

**Abbreviations**: L, left; R, right; MJS, Multidimensional Jealousy Scale; AQ, Buss-Perry aggression Questionnaire.

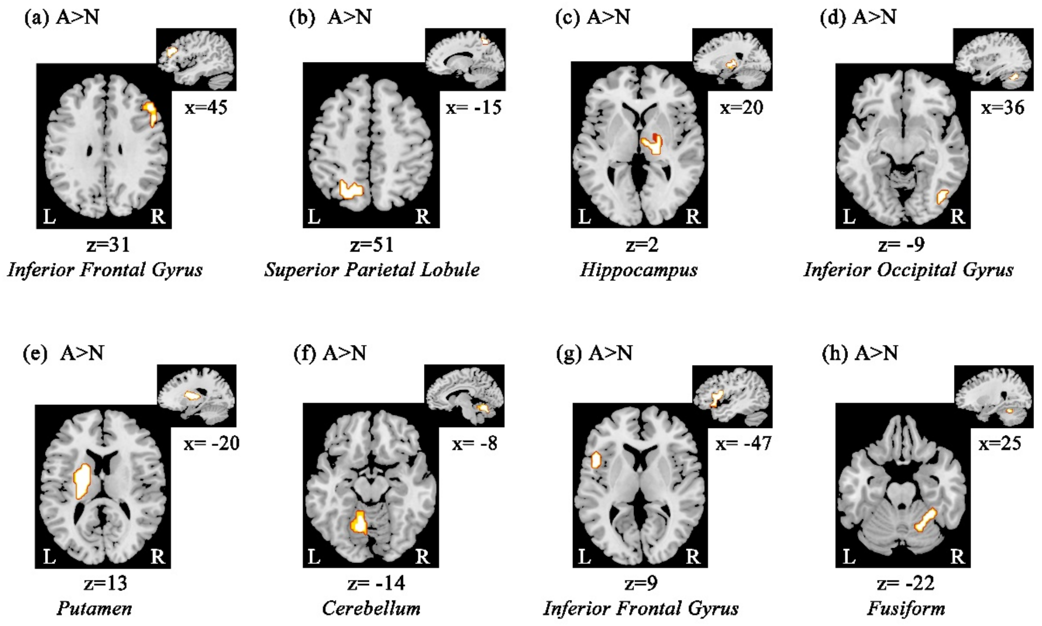


**Figure S1. Whole brain analysis *(a) - (h)*** Regions showing increased activation with higher jealousy traits (as assessed by the MJS) during processing angry facial expressions relative to neutral faces, with gender, intensity rating of angry faces and AQ as covariates. Findings are displayed at $p < 0.05$ cluster-level FWE correction with a cluster-forming threshold $p < .001$. Coordinates of peak voxels (x/y/z) are given in Montreal Neurological Institute (MNI) space.

**Abbreviations**: A>N, contrasts angry > neutral; L, left; R, right; MJS, Multidimensional Jealousy Scale; AQ, Buss-Perry aggression Questionnaire.

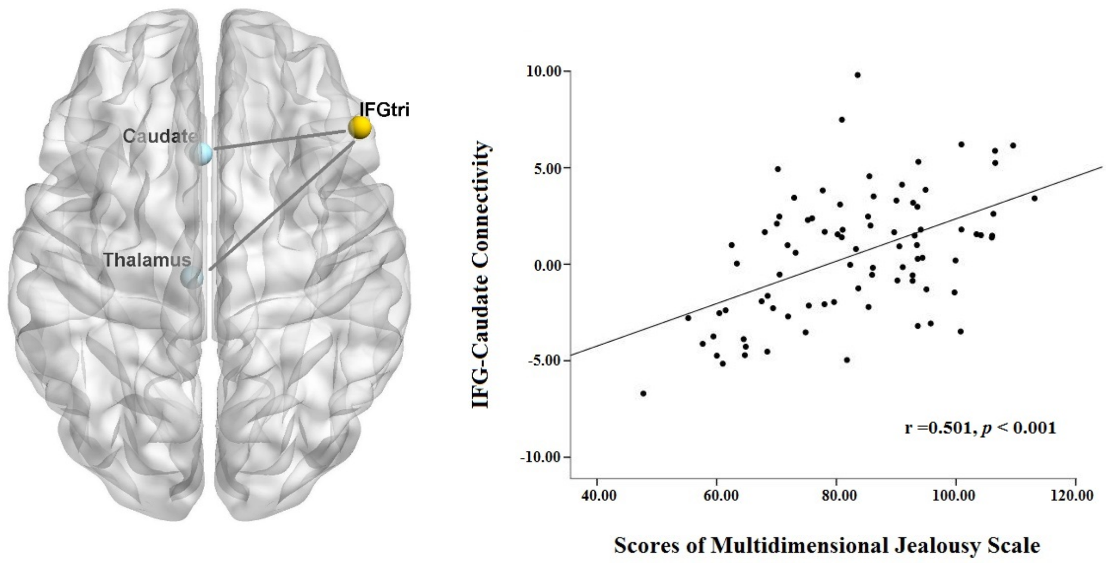

**Figure S2. Associations on the task-based network level. (a)** *Right Triangular Inferior Frontal Gyrus – IFGtri(x/y/z: 51/29/26) as seed region*. Right IFGtri functional connectivity with the left caudate (x/y/z: -3/20/5), and the left thalamus(x/y/z: -6/-22/8) were positively correlated with trait jealousy (MJS) during processing of angry relative to neutral faces. **(b)** Scatter visualization of the association between trait jealousy and IFGtri – left caudate coupling using extracted parameter estimates. Results were significant at $p < 0.05$ cluster-level FWE correction with an initial cluster-forming threshold $p < .001$. Coordinates of peak voxels (x/y/z) are given in Montreal Neurological Institute space.
